# Vermont: a multi-perspective visual interactive platform for mutational analysis

**DOI:** 10.1101/166348

**Authors:** Alexandre V Fassio, Pedro M Martins, Samuel da S Guimarães, Sócrates S A Junior, Vagner S Ribeiro, Raquel C de Melo-Minardi, Sabrina de A Silveira

## Abstract

**Background:** A huge amount of data about genomes and sequence variation is available and continues to grow on a large scale, which makes experimentally characterizing these mutations infeasible regarding disease association and effects on protein structure and function. Therefore, reliable computational approaches are needed to support the understanding of mutations and their impacts. Here, we present VERMONT 2.0, a visual interactive platform that combines sequence and structural parameters with interactive visualizations to make the impact of protein point mutations more understandable.

**Results:** We aimed to contribute a novel visual analytics oriented method to analyze and gain insight on the impact of protein point mutations. To assess the ability of VERMONT to do this, we visually examined a set of mutations that were experimentally characterized to determine if VERMONT could identify damaging mutations and why they can be considered so.

**Conclusions:** VERMONT allowed us to understand mutations by interpreting position-specific structural and physicochemical properties. Additionally, we note some specific positions we believe have an impact on protein function/structure in the case of mutation.

## Background

According to the International HapMap Project [1], there are approximately 10 million common single-nucleotide polymorphisms (SNPs); whereas, in accordance with the 1,000 Genomes Project Consortium, the difference between the genome of an individual selected at random and the reference genome is approximately 10,000 non-synonymous SNP (nsSNP) sites [2]. SNPs represent more than half of all the disease-associated variations in the Human Gene Mutation Database (HGMD) [3].

The sequence variation in a genome is a complex phenomenon. A huge amount of data involving genomes and especially sequence variation is available and continues to grow on a large scale. This makes experimentally characterizing these variations in terms of disease association and effects on protein structure and function infeasible. Therefore, reliable computational approaches are needed to support the understanding of mutations and their impacts.

Over the past two decades, several computational methods have been proposed to understand and predict the influence of mutations in protein structure and function based on different evolutionary and physicochemical data. Two recent reviews gave a panorama of such tools by discussing some representative cases, with some overlap [2, 4]. We did not aim to develop an exhaustive list of such methods because we believe this was already done well in the mentioned reviews. In this paper, we comment on some recent strategies that have been proposed to understand and predict the impact of mutations on protein structure and function based on different perspectives.

Worth and colleagues proposed in [5] the web server Site Directed Mutator (SDM) [6], which uses a statistical potential energy function to predict the effect of SNPs on the stability of proteins based on environment-specific amino acid substitution frequencies within homologous protein families.

In [7], Pires and others introduced mCSM, which encodes distance patterns between atoms to represent protein residue environments as graphs, where nodes are the atoms and the edges are the physicochemical interactions established among them. From these graphs, distance patterns are extracted and summarized in a structural signature that is used as evidence to train predictive models.

Also based on graphs, in [8] Giollo and colleagues proposed NeEMO, a non-linear neural network model for the prediction of stability changes upon mutations based on residue interaction networks (RINs). RINs are a graph description of protein structures where nodes represent amino acids and edges represent different types of physicochemical interactions.

Laimer and others, in turn, proposed multi-agent stability prediction upon point mutations (MAESTRO) [9]. The method combines multiple linear regression, a neural network approach and support vector machine (SVM) with a multi-agent method to predict protein structure stability mainly based on ΔΔ*G*. In [10], the authors present MAESTROweb, a web interface for MAESTRO (which is a standalone software).

A predictor of the Impact of Non-synonymous variations on Protein Stability (INPS) was introduced in [11]. This method computes the ΔΔ*G* values of protein variants without relying on the protein structure, taking advantage of the fact that the number of available sequences is much higher than the number of structures. In [12], the authors presented INPS-Multi Descriptor (MD), which complements INPS with a new predictor (INPS-3D) that exploits descriptors derived from the protein structure.

iStable, proposed in [13], integrates I-Mutant2.0 [14], MUPRO [15], AUTO-MUTE [16], PoPMuSiC2.0 [17], and CUPSAT [18] through SVM to predict protein stability changes upon single amino acid residue mutations, and it performs better than any single method alone.

DUET, presented in [19], combines mCSM [7] and SDM [5] to predict the effects of missense mutations by consolidating the results of both methods in an optimized predictor through SVM trained with Sequential Minimal Optimization [20].

Despite several strategies being proposed to predict the impact of mutations, none of them alone has been proven to be accurate in all scenarios where mutation impact is investigated [19]. Under these circumstances, a strategy that has gained attention is combining methods based on different paradigms and protein structural properties for the purpose of reaching a consensus on the understanding of mutation impacts. iStable and DUET are examples of such methods. Another inconvenience regarding the methods that are widely used in the study of a mutation’s impact is the lack of interpretability.

Authors from the mentioned works on protein mutations and from the reviews [2, 4] note common directions that can be explored to develop strategies with more accurate predictions. Some notable guidelines are the use of consensus approaches that integrates various methods; the development of user-friendly tools; and the use of relevant features to better describe the properties of mutations, such as those based on sequence, structure and database annotation. In line with these directions, this article proposes ViewER MutatiON Tool (VERMONT) 2.0, a visual interactive platform that integrates sequence and structural parameters such as intramolecular interactions, solvent accessibility, and topological properties, coupled with powerful interactive visualizations to make the impact of protein point mutations more understandable. VERMONT is visual analytics oriented, so it allows domain specialists to analyze and make sense of many structural properties for gaining insights into the impact of point mutations.

The first version of VERMONT [21] was presented in Biovis Contest in 2013 to analyze data from a functionally defective triosephosphate isomerase (dTIM) and its S. cerevisiae parent (scTIM) based on a dataset of proteins of the same family. The main goal was to point out mutations that have an impact on function and suggest how the function could be rescued. At that time, VERMONT was populated only with the contest data, and it was not possible to analyze mutations in proteins other than dTIM.

Due to the positive feedback of VERMONT, which received the *Biology Experts Pick* award, we decided to implement a whole new VERMONT 2.0 from scratch. Now, the tool takes as input any protein structure in PDB file format. The input module automatically searches the Protein Data Bank [22] for similar structures given an entry informed by the user and according to a desired similarity threshold, or the user can enter a list of PDB entries. Then, VERMONT proceeds to the necessary computations and notifies the user when the analysis has been completed. Furthermore, new interaction graphs and protein molecular structural visualizations were included to potentialize the analysis of specialists. We also coupled in our platform, the FoldX tool [23], which predicts the impact of a mutation through the calculation of Gibbs free energy change (ΔΔ*G*) to complement visual parameters displayed in VERMONT, supporting users on the selection of harmful mutations.

## Methods

In this section, we detail the VERMONT platform by describing problem modeling, its functionalities and interactive visualizations organized by modules. A summary of the VERMONT analysis process is presented in Figure 1.

**Figure 1.**
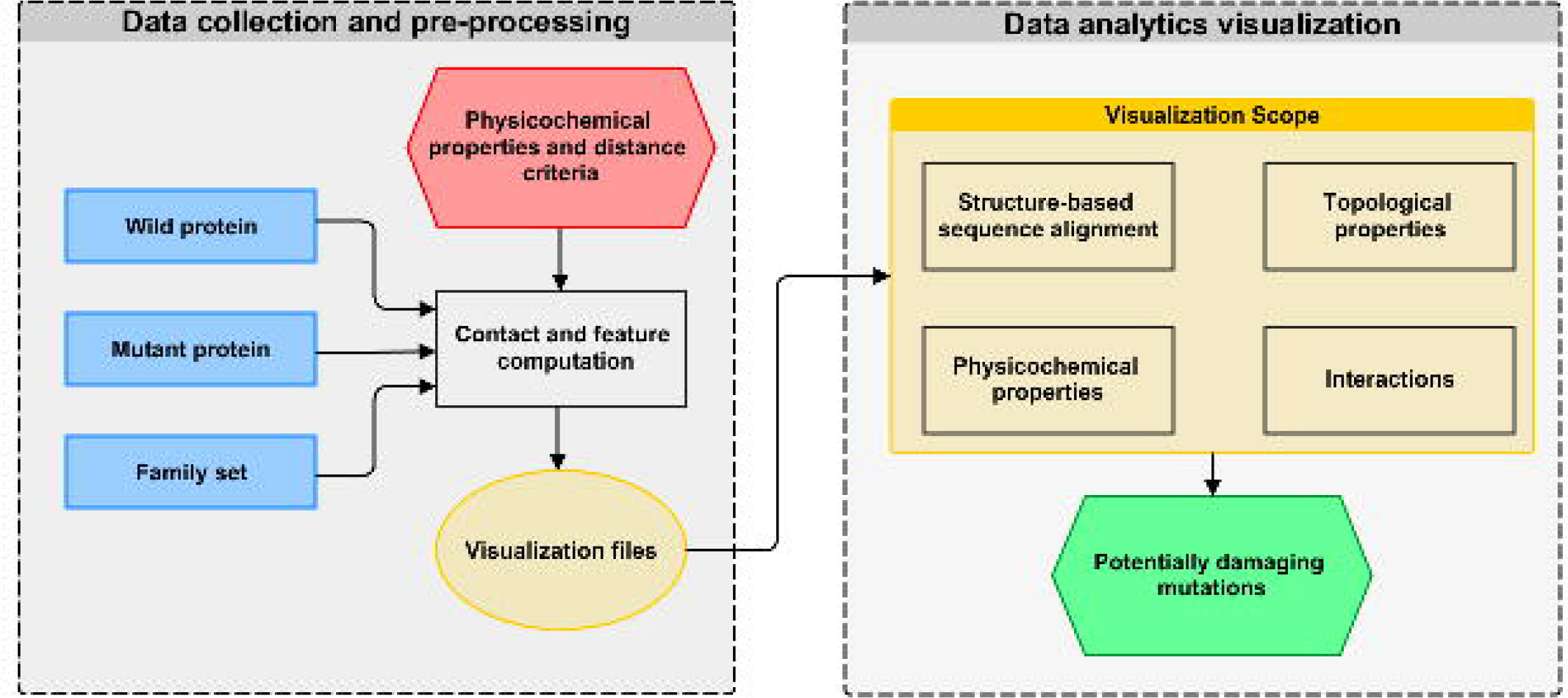
VERMONT scheme. The first step of VERMONT is data collection and preprocessing of wild type and mutant proteins, as well as the family set, for computation of structure-based alignment, accessibility, topological properties, and interactions data. Next, in the visualization modules, domain specialists can explore and interpret all computed features to note potentially damaging mutations.

Given a dataset, we compute a variety of sequence and structural parameters for each residue. We were interested in visually representing these parameters in a way that they can be examined to detect relevant similarities and differences as well as trends and exceptions, which constitutes a multivariate visualization problem.

A first task that domain specialists perform to identify similarities and differences among a set of proteins is a sequence alignment, which shows each protein sequence in a row and equivalent residues in the same column. In addition, residues are usually colored according to a color scheme that associates residues with similar physicochemical properties to the same color, helping to identify conservation in a particular column.

To take advantage of a visual representation that domain specialists are very familiar with, we used the multiple sequence alignment visualization as the basis for our platform. In addition to displaying sequence alignment, we include an intramolecular interaction network, solvent accessibility, physicochemical properties and complex network topological parameters in this basic sequence alignment visualization.

### Visualized attributes computation methods

Each structure was modeled as a graph to study the network of intramolecular interactions and analyze its topological properties from a complex network perspective. We computed interatomic contacts using Delaunay triangulation [24], which is a geometric and cutoff independent approach where edges represent interatomic interactions, excluding occluded contacts. Contact computation was performed using the CGAL [25] library, version 3.3.1.

For each chain of a particular PDB id, we constructed an atomic level contact graph where nodes represent atoms and edges represent interactions among them. Nodes are labeled according to their physicochemical properties as positive, negative, hydrogen bond donor, hydrogen bond acceptor, aromatic, hydrophobic and cysteine based on our previous works [26, 27], which were, in turn, derived from [28]. Edges are labeled according to interatomic interactions and distance criteria such as hydrogen bond, repulsive, salt bridge, aromatic, hydrophobic and disulfide bridge based on [29]. The interactions were then mapped to residue level.

These graphs, which represent protein structures, are the basis for the *Interactions* and *Topological properties* modules of VERMONT. Table 1 provides the distance criteria and atom labels for each interaction type.

**Table 1.**
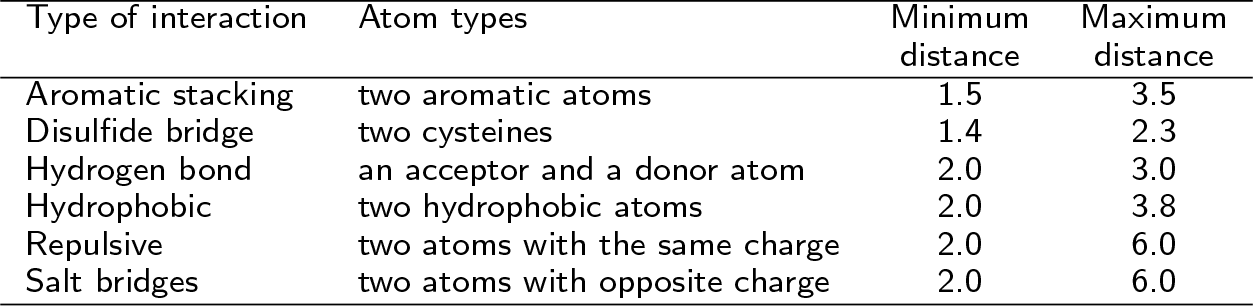
Distance criteria (in Å) and atom types for each interaction.

| Type of interaction | Atom types | Minimum distance | Maximum distance |
| --- | --- | --- | --- |
| Aromatic stacking | two aromatic atoms | 1.5 | 3.5 |
| Disulfide bridge | two cysteines | 1.4 | 2.3 |
| Hydrogen bond | an acceptor and a donor atom | 2.0 | 3.0 |
| Hydrophobic | two hydrophobic atoms | 2.0 | 3.8 |
| Repulsive | two atoms with the same charge | 2.0 | 6.0 |
| Salt bridges | two atoms with opposite charge | 2.0 | 6.0 |

### Common features of VERMONT modules

Next, we describe some features that are common to more than one visualization module in VERMONT.

**Selection buttons:** These show All alignment colors, only Mutant columns or only CSA columns (catalytic site residues), taking as reference the wild protein.

**Color filters:** Residues can be selected to be colored according to each group of the color scheme or not.

**Zoom control:** This control is provided by a slider so data can be visualized in small values of zoom to give a panorama of the structural alignment, which helps in detecting general trends and exceptions, such as highly conserved columns and columns that are not at all conserved. Higher values for zoom allows viewing details about the residues and the alignment. To have details on demand about a particular position, the user needs to hover the mouse over it, which opens a pop-up with an alignment position, real position (the position on the PDB sequence), residue and PDBid.chain.

**Frequent residues by position:** This highlights columns that have the selected percentage of conservation.

**Select N columns by average:** Given a topological or physicochemical property, it highlights the N first columns with higher (top-down) or lower (bottom-up) average values.

**Energy variation prediction:** The effect on protein stability is evaluated by calculating the ΔΔ*G* for each mutation using FoldX tool. In the visualization module, these values are highlighted by color-coded rectangles that vary from red (highly destabilizing) to gray (neutral) to blue (highly stabilizing), while values not calculated are colored in yellow. Moreover, detailed information can be accessed by hovering the mouse over a mutation position on the mutant sequence. Details about ΔΔ*G* range are in *Additional file 1*.

**Sequence logo:** This is positioned below the alignment panel, it represents the sequence conservation for each column by depicting the consensus sequence as well as the diversity of each position.

### Input module

The VERMONT input module is shown in Figure 1 from the Additional file 1, and it takes three basic parameters:

- The structure of a wild protein: a PDB identifier and chain, which we will call PDBid.chain from now on;
- The sequence of the mutant protein: a sequence in FASTA file format that represents the same wild protein after mutations;
- A set of proteins, which we call a family: a set of protein structures (PDBid.chain) similar to the wild protein. In this case, the user has two options: (i) select an alignment method (BLAST, FASTA, PSI-BLAST) and a similarity threshold to allow VERMONT to search on PDB for the set of proteins; or (ii) inform a set of structures the user considers similar to the wild protein.

Additionally, users can receive an email to be notified when the server finishes data processing.

### Structure-based sequence alignment module

A structure-based sequence alignment of each protein from the family set against the wild protein is performed in a pairwise manner using Multiprot [30]. To represent this alignment, we used multiple sequence alignment visualization, a kind of visualization biologists use to analyze and visualize. This visualization is the basis of our strategy, and an example of the *Structure-based sequence alignment* module is provided in Figure 2. Sequences from the family set are stacked using the wild protein sequence, on the top, as a reference. The sequence of the mutant protein is then positioned above the wild protein. Each row and column represent a protein chain and a correspondent position in the alignment, respectively. Each residue is colored according to its physicochemical properties, and similar rows are organized next to each other using the clustering algorithm Expectation Maximization (EM). The coloring and clustering helps to identify conservations and exceptions in columns.

**Figure 2.**
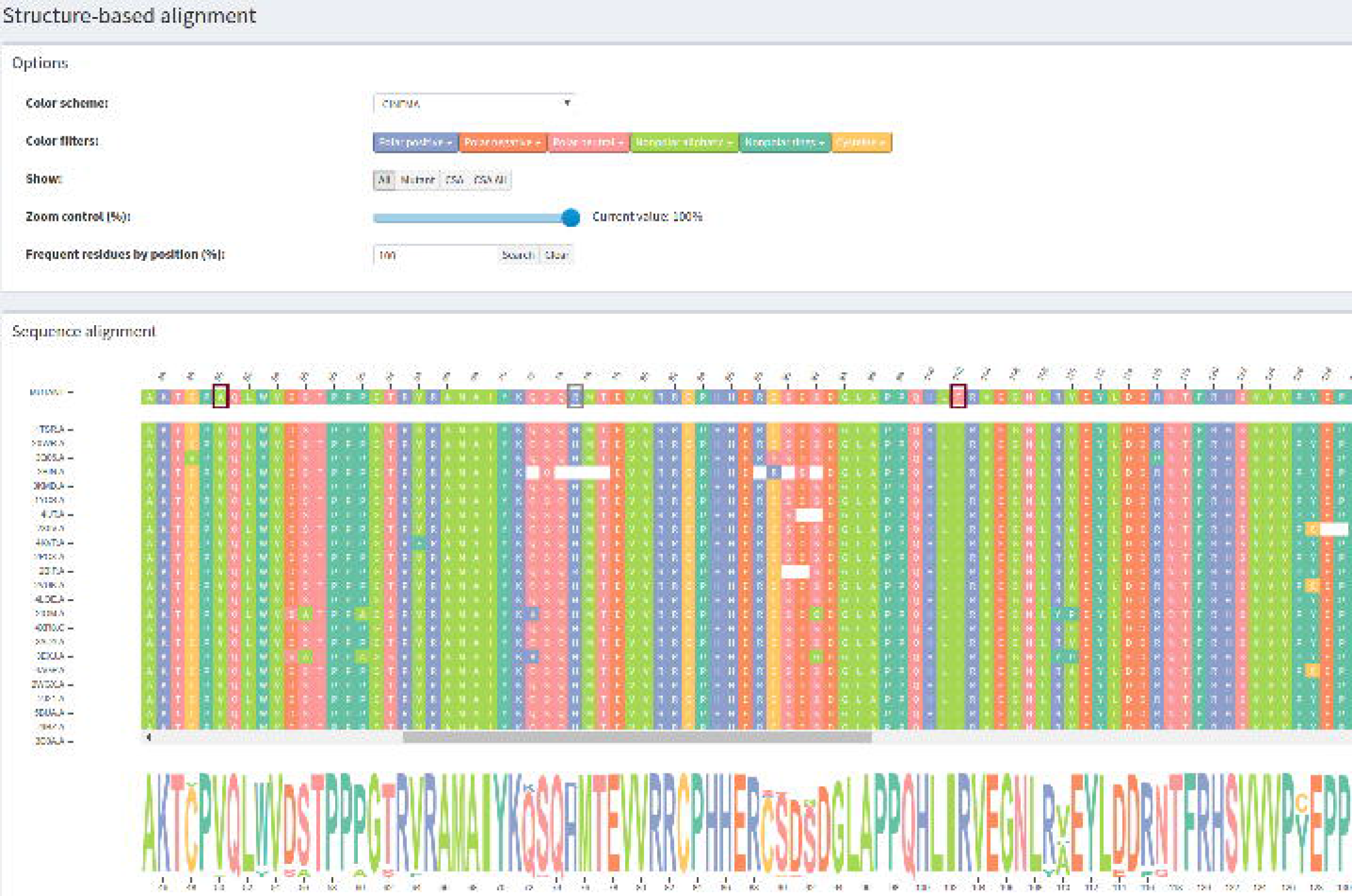
Structure-based sequence alignment module. These data are from wild type protein p53 (1TSR.A) and its variants were selected using 70% identity and the PSI-BLAST alignment method.

Three color schemes are provided for protein residues:

- CINEMA: distinguishes among 6 groups, which are polar positive in blue (H, K, R), polar negative in orange (D, E), polar neutral in pink (N, Q, S, T), nonpolar aliphatic in light green (A, G, I, L, M, V), nonpolar rings in dark green (F, P, W, Y) and cysteine in yellow (C).
- CLUSTAL: segments residues in 4 groups that are (G, P, S, T) in yellow, (H, K, R) in orange and (F, W, Y) in blue and (I, L, M, V) in light green.
- LESK: uses 5 groups, which are small nonpolar in orange (G, A, S, T), hydrophobic in green (C, V, I, L, P, F, Y, M, W), polar in magenta (N, Q, H), negatively charged in red (D, E) and positively charged in blue (K, R).

After selecting a color scheme, there are some features to help users analyze and make sense of the data that are common in VERMONT modules, so we describe them separately in the Section Common features of VERMONT modules.

### Interactions module

The intramolecular interactions of each structure are represented as a graph as detailed in the Section *Visualized attribute computation methods*. However, it is not trivial to identify and grasp conserved patterns in protein interactions by visually inspecting graphs. Thus, we devised a 2D representation of intramolecular interactions that gives a panorama of the intramolecular network, delineating the conserved columns for the whole *family* dataset at once. An example of the *Interactions* module is provided in Figure 3. In Figure 3(A), we show all interactions at once, while we show the interactions for a selected column in Figure 3(B).

**Figure 3.**
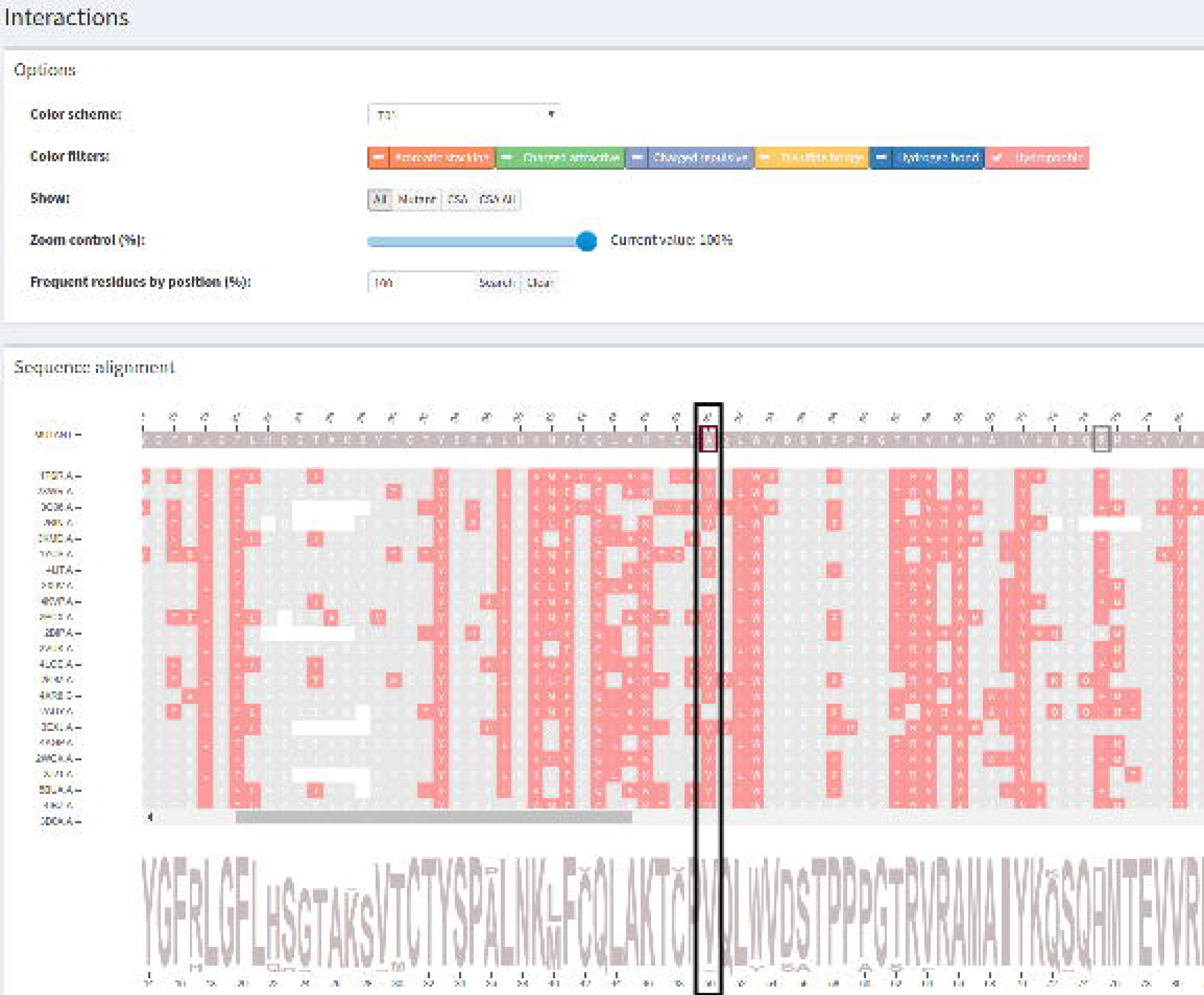
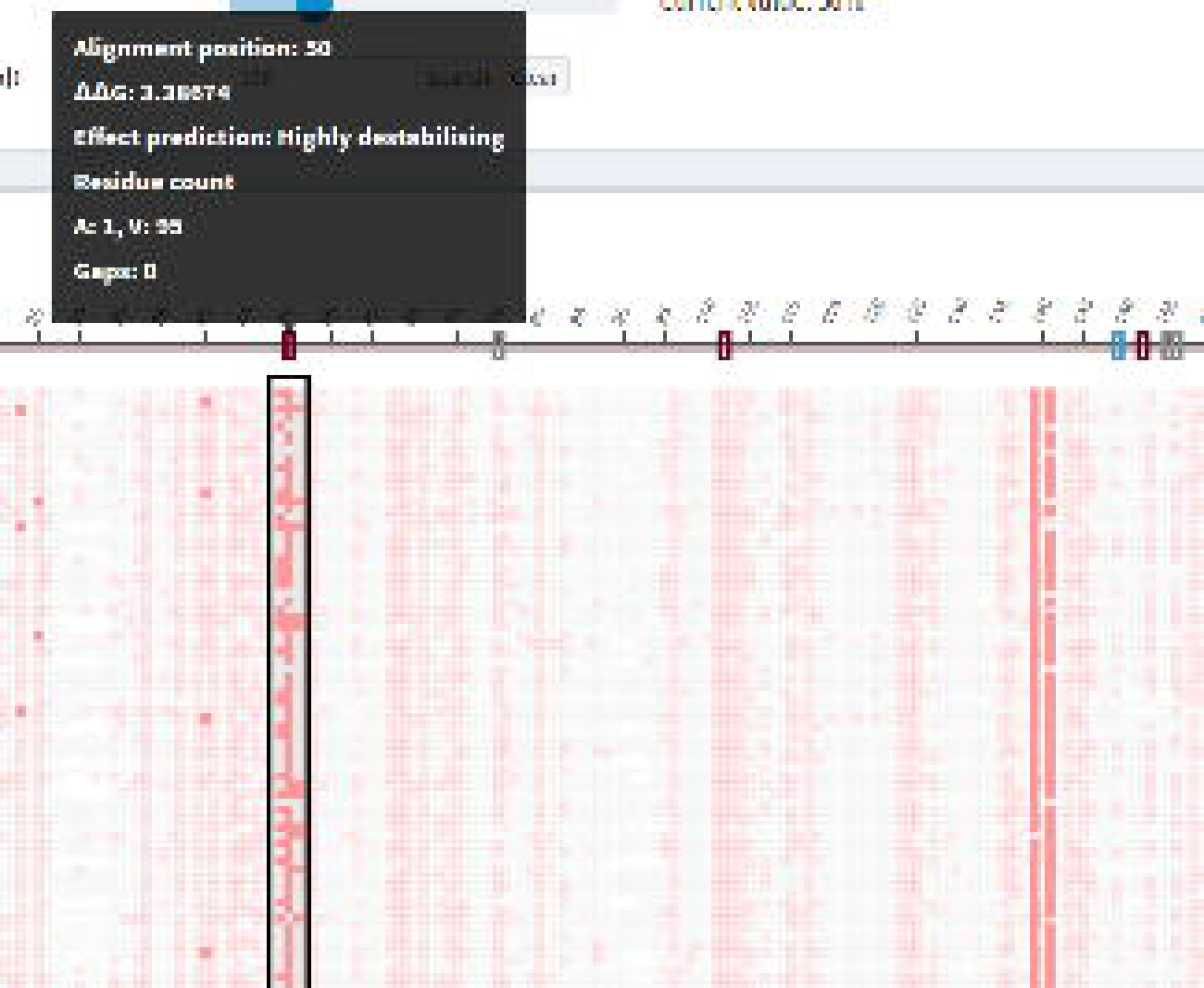
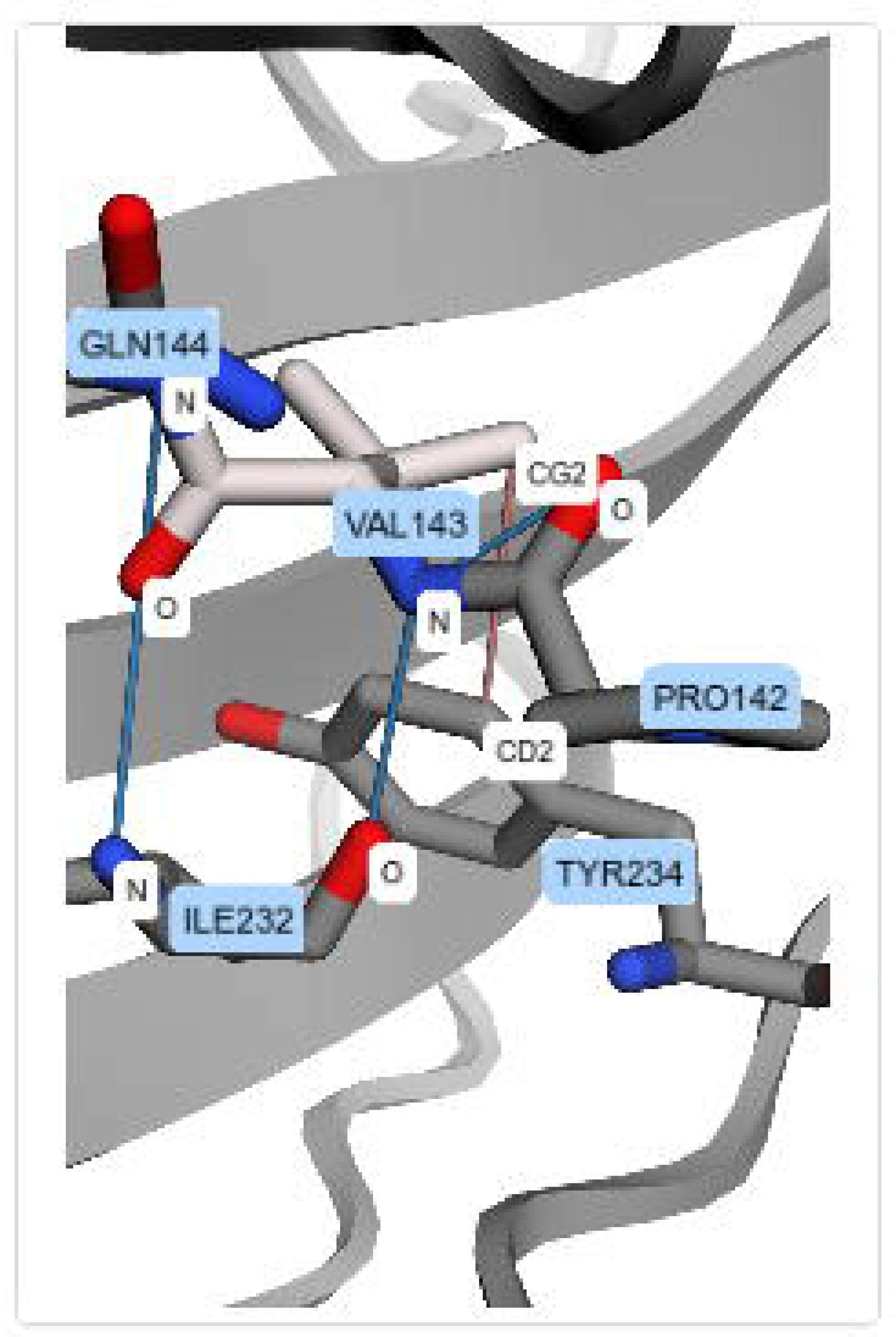
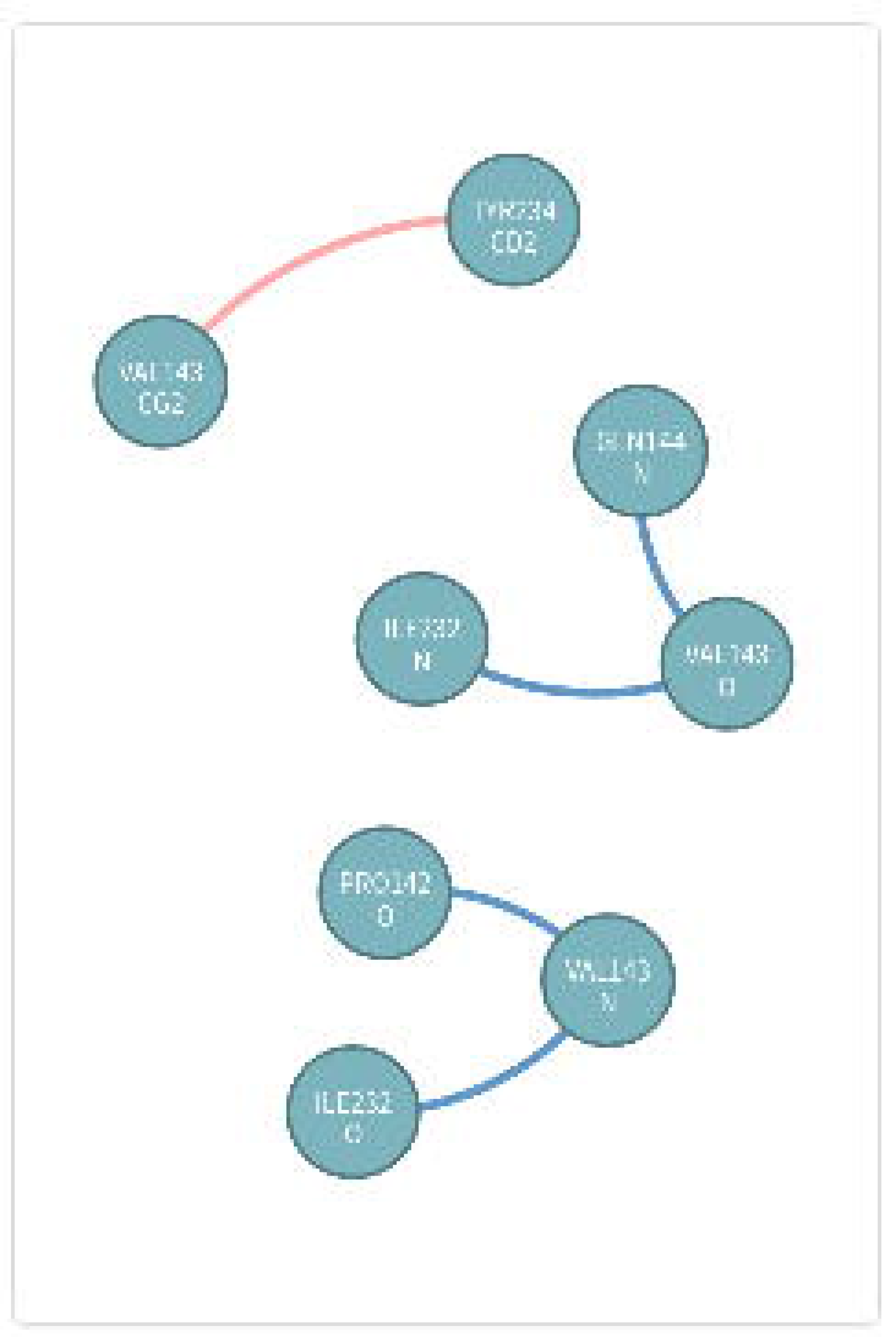
Interactions module. The displayed data are from wild type protein p53 (1TSR.A) and its variants were selected using 70% identity and the PSI-BLAST alignment method. (A) All residues that establish hydrophobic interactions are colored in rose. The alignment position 50, which corresponds to mutation Val143Ala, was highlighted. (B) Hydrophobic interactions for mutation Val143Ala. The alignment position (column) 50 was selected to show only its hydrophobic interactions. The zoom of 30% was used to display short and long distance interactions. (C) 3D molecular representation of interactions for Val143 from protein p53 (1TSR.A). (D) Graphs (2D schematic representation) of p53 (1TSR.A) interactions for residue Val143.

The multiple sequence alignment visualization, which is the basis of our tool, is used to represent the interactions. Each residue is colored according to the interaction it establishes. If a residue establishes more than one interaction, it is colored in gray. Hence, VERMONT provides a general view of the interactions, delineating the conserved columns. Additionally, one can select a specific column to inspect its contacts, which points out specific patterns on the contacts of a correspondent position in the alignment. The user can choose to show or hide each type of interaction in the sequence alignment panel.

By clicking on a particular position (a residue) in the sequence alignment visualization, VERMONT shows the *Interaction viewer*. In this module, the interactions established by the selected residue are depicted as a 3D molecular representation of the protein (Figure 3(C)) and as a 2D schematic representation in the form of a graph (Figure 3(D)), which allows users to make sense of these interactions in the context of protein structures.

Some interactions involve residues that are close to each other in the sequence space while others involve residues that are distant. To support users on the visualization and analysis of both long and short range contacts, we have a *zoom control* to provide a general view of the interactions, maintaining long and short range contacts on the same screen by using low values for zoom. Contact details can be obtained by using high values for zoom, hovering the mouse over each residue to see more information or by clicking on a specific residue to see its interactions in 3D and in 2D representations.

### Topological properties module

Complex networks are graphs whose connections between nodes are neither purely regular nor purely random. Most real-world graphs, such as for protein–protein interactions or social or gene-regulatory networks, are complex [31].

In VERMONT, three common complex network centrality measures were computed for each residue; that is, each node from each graph that represents a protein structure. These metrics were computed using the iGraph [32] package, version 1.0.1. Here, we briefly describe them. In the *Additional file 1*, we describe these metrics in detail and some of their uses and meanings in biology. Figure 4 shows the topological properties panel.

- Degree: the degree of a vertex in a graph is the number of edges connected to it.
- Betweenness: the extent to which a vertex lies on paths between other vertices.
- Closeness: the mean distance from a vertex to all other vertices.

**Figure 4.**
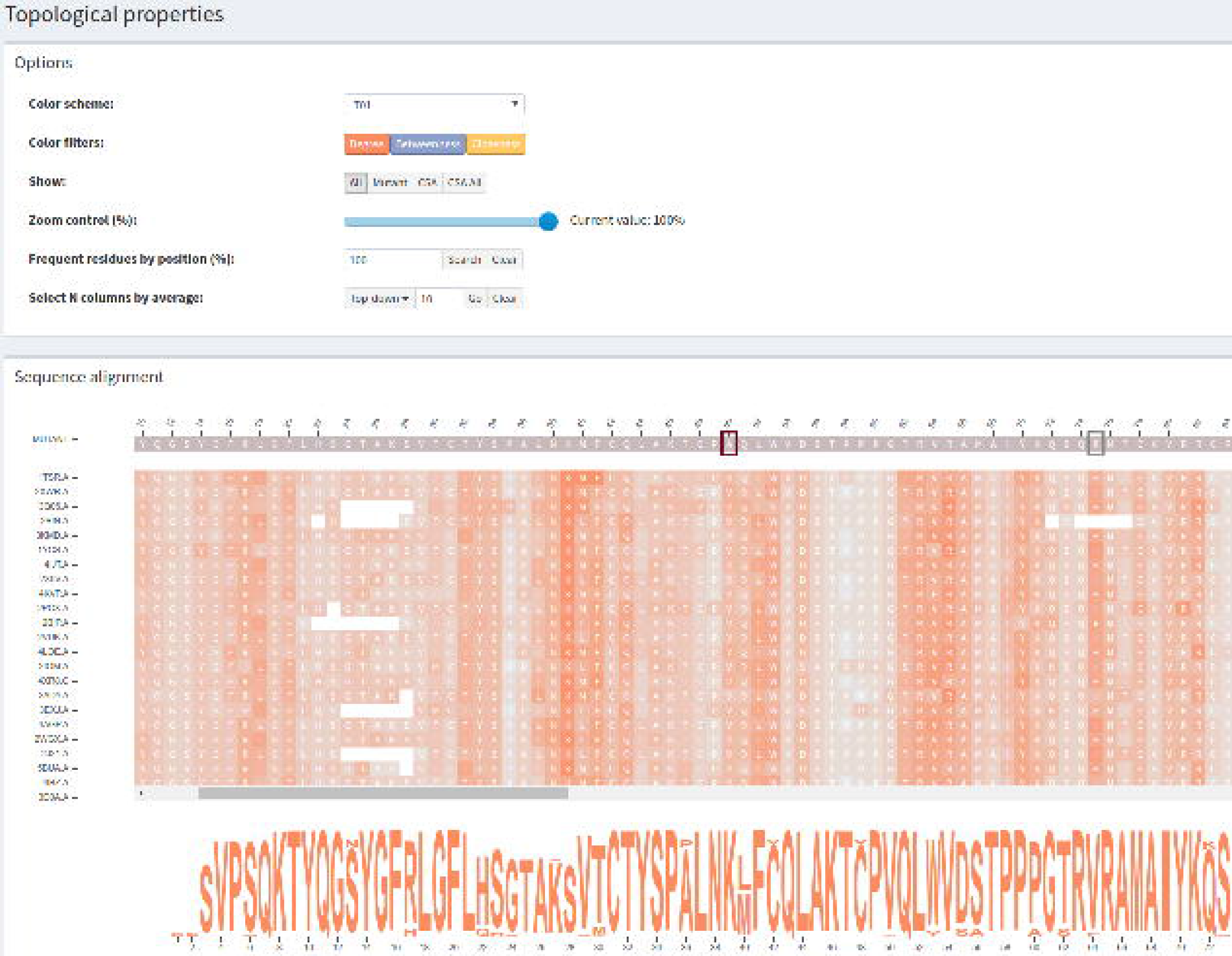
Topological properties module. The degree centrality measure for wild type protein p53 (1TSR.A) and its variants selected using 70% identity and the PSI-BLAST alignment method. Network centralities are displayed in a heatmap based on the sequence alignment visualization.

These network topological properties are displayed in VERMONT using a heatmap constructed based on the multiple sequence alignment visualization. Each measure (degree in orange, betweenness in blue and closeness in yellow) is shown on a specific heatmap panel. Individual residues contained in the alignment visualization are represented as color intensities.

This heatmap representation supports users by detecting relevant residues/positions in the alignment from the complex network perspective. Columns with high values of topological properties are shown in a dark shade of the selected color and columns with low values are shown in light shades. As a column corresponds to a specific position in the alignment, columns that exhibit a trend should be further investigated.

### Accessibility module

Solvent accessibilities were computed through Lee and Richards algorithm [33] using the software Naccess. This software calculates the accessible area by rolling a probe of a given radius (typically 1.4 °A, as it is the water radius) around the Van der Waal’s surface of the protein. The path traced out by the probe center is the accessible surface. Figure 5 shows the *Accessibility* module.

**Figure 5.**
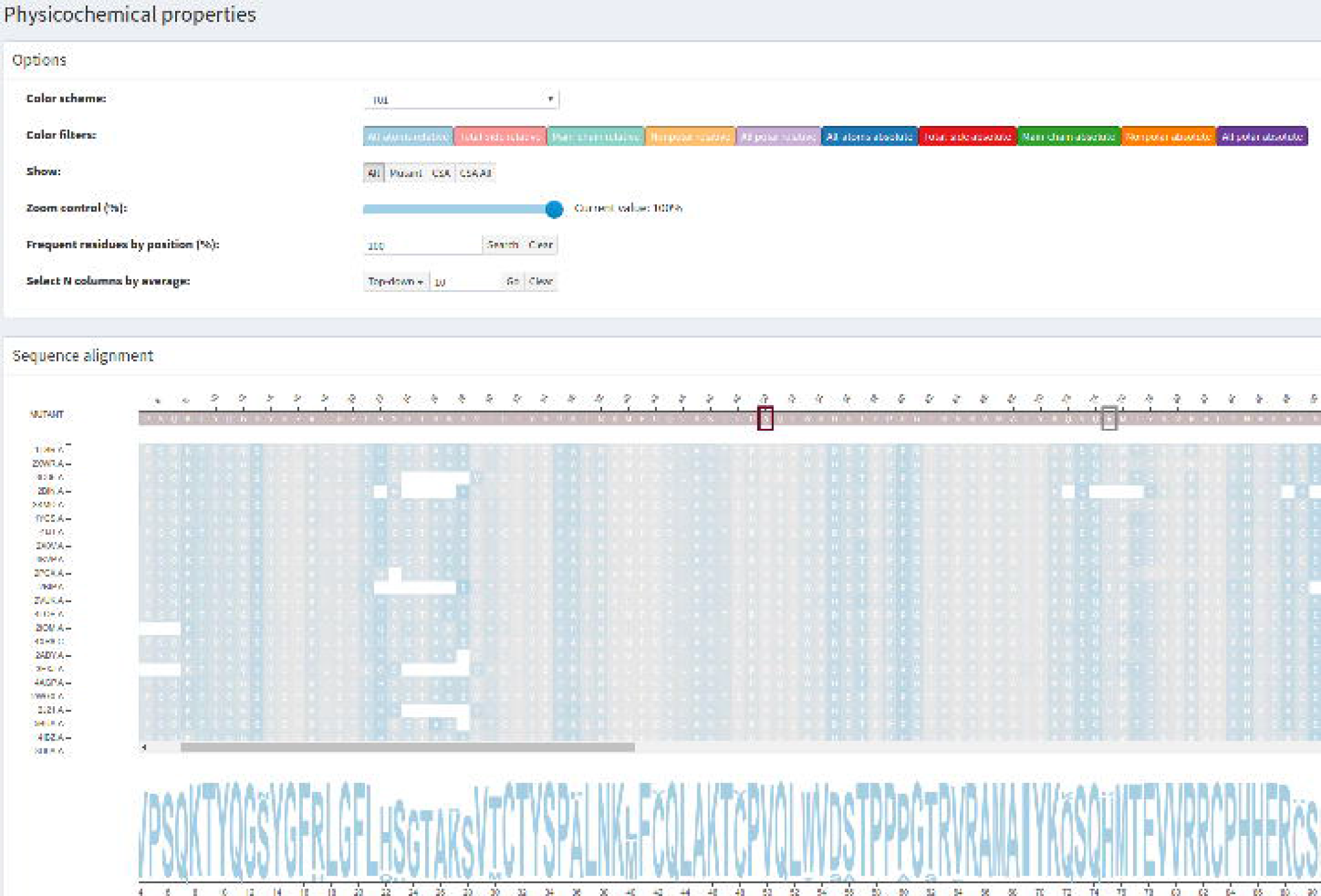
Accessibility module. Accessibilities computed using Naccess are displayed in a heatmap based on the sequence alignment visualization. The all-atoms relative accessibility that is displayed is for wild type protein p53 (1TSR.A), using 70% identity and the PSI-BLAST alignment method.

Hydrophobic interactions are important forces in initializing protein folding and stabilizing 3D structures of proteins. Hydrophobicity and the packing of hydrophobes in the hydrophobic core of a protein can affect protein stability [34]. In globular proteins, the hydrophobic (apolar) residues are bounded towards the protein core, forming hydrophobic cores, whereas hydrophilic (polar) residues are more exposed to solvent. This hydrophobic packing in the protein core tends to be conserved in protein families. Thus, we believe a mutation in the protein core is more likely to be destabilizing than a mutation on the protein surface, with some exceptions as, for instance, mutations in the binding site and the active site.

We combined a multiple sequence alignment visualization with a heatmap to display accessibilities. We provide one heatmap for each accessibility computed using Naccess, which are *all-atoms relative*, *total-side relative*, *main-chain relative*, *nonpolar relative*, *all polar relative*, *all-atoms absolute*, *total-side absolute*, *main-chain absolute*, *nonpolar absolute* and *all polar absolute*. Each alignment position, which corresponds to a residue, is associated with a color intensity. The higher the value, the more intense the color. The lower the value, the less intense the color. This heatmap allows users to detect conserved columns (correspondent positions) in the alignment, which means columns that have high or low values of accessibility.

## Results and discussion

To assess the ability of VERMONT to support domain specialists when analyzing a large amount of structural properties to gain insights on the impact of point mutations, selecting those that are potentially damaging for further investigation, we performed a use case in which we selected a classical mutation dataset from Bongo [35], which has been used in many subsequent studies as [7, 19, 8, 11, 12]. We visually examine the mutations by integrating the sequence conservation, intramolecular interaction network, solvent accessibility, physicochemical properties and complex network topological parameters to gain insights into the impact of mutations. Additionally, we note a few mutations that could be potentially damaging according to VERMONT.

### Use case

The p53 gene encodes a transcription factor with multiple, anti-proliferative functions activated in response to several forms of cellular stress. The core domain of tumor suppressor protein, p53, is responsible for approximately 50% of the mutations that lead to human cancers [36]. Eight disease-associated mutations in the p53 core domain that were analyzed experimentally by Fersht and co-workers [37, 38] were used in this use case. In Table 2, we provide these eight mutations. Next, we describe how two of these mutations, Arg273His and Ile195Thr, could be visually analyzed as illustrative cases using VERMONT. The other six mutations are described in the *Additional file 1* due to space limitations. In this analysis, we considered the all-atoms relative accessibility. We worked with relative accessibilities as they express the accessible surface as a percentage of that observed in an Ala-X-Ala tripeptide.

**Table 2.**
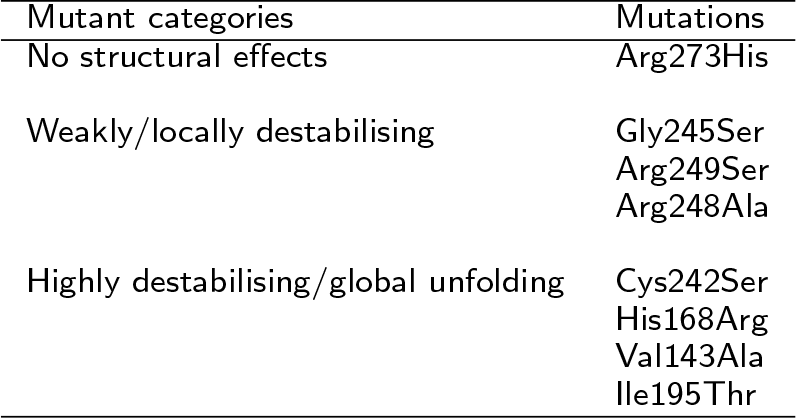
Mutations (nsSNPs) in the p53 (PDBid 1TSR) core domain that were experimentally characterized.

The input parameters used in VERMONT were (i) PDB id 1TSR.A as the wild protein; (ii) the mutant fasta file, generated by manually changing original residues in the 1TSR.A fasta file by those that are the result of mutations; (iii) PSI-BLAST as the alignment method; and (iv) 70% identity. The results are available to be explored and analyzed in VERMONT. A summary of the results obtained for accessibility, topological properties, and interactions are presented in Tables 3 and 4.

**Table 3.** Summary of accessibility and topological parameters computed by VERMONT for mutations (nsSNPs) in the p53 (PDBid 1TSR) core domain that were experimentally characterized.

|  | Accessibility |  | Degree |  | Closeness (E-04) |  | Betweenness |  |
| --- | --- | --- | --- | --- | --- | --- | --- | --- |
| | $\mu$ | $\sigma$ | $\mu$ | $\sigma$ | $\mu$ | $\sigma$ | $\mu$ | $\sigma$ |
| Arg273His | 30.60 | 6.53 | 6.68 | 1.10 | 9.69 | 1.64 | 1487.24 | 567.77 |
| Gly245Ser | 8.81 | 6.97 | 2.91 | 0.60 | 7.77 | 1.38 | 225.21 | 113.30 |
| Arg249Ser | 15.46 | 4.38 | 5.30 | 1.13 | 8.66 | 1.48 | 635.93 | 286.51 |
| Arg248Ala | 72.98 | 8.74 | 3.13 | 0.47 | 8.22 | 1.41 | 335.64 | 263.27 |
| Cys242Ser | 13.82 | 7.24 | 3.31 | 0.73 | 7.37 | 1.18 | 200.86 | 130.92 |
| His168Arg | 26.46 | 4.59 | 5.86 | 0.83 | 8.57 | 1.42 | 799.43 | 267.57 |
| Val143Ala | 0.31 | 0.55 | 4.00 | 0.46 | 9.55 | 1.69 | 642.42 | 407.30 |
| Ile195Thr | 5.76 | 0.70 | 3.88 | 0.70 | 10.51 | 2.06 | 1193.47 | 455.24 |

**Table 4.** Summary of interactions computed by VERMONT for mutations (nsSNPs) in the p53 (PDBid 1TSR) core domain that were experimentally characterized.

|  | Aromatic stacking |  | Charged attractive |  | Charged repulsive |  | Disulfide bridge |  | Hydrogen bond |  | Hydrophobic |  |
| --- | --- | --- | --- | --- | --- | --- | --- | --- | --- | --- | --- | --- |
| | $\mu$ | $\sigma$ | $\mu$ | $\sigma$ | $\mu$ | $\sigma$ | $\mu$ | $\sigma$ | $\mu$ | $\sigma$ | $\mu$ | $\sigma$ |
| Arg273His | 1.00 | 0.00 | 6.79 | 1.51 | 1.72 | 1.00 | 0.00 | 0.00 | 4.49 | 0.85 | 1.00 | 0.00 |
| Gly245Ser | 0.00 | 0.00 | 0.00 | 0.00 | 0.00 | 0.00 | 0.00 | 0.00 | 2.90 | 0.61 | 1.00 | 0.00 |
| Arg249Ser | 0.00 | 0.00 | 3.56 | 0.68 | 1.20 | 0.76 | 0.00 | 0.00 | 5.09 | 1.46 | 1.44 | 0.50 |
| Arg248Ala | 0.00 | 0.00 | 1.00 | 0.00 | 2.13 | 0.88 | 0.00 | 0.00 | 3.04 | 0.50 | 0.00 | 0.00 |
| Cys242Ser | 0.00 | 0.00 | 0.00 | 0.00 | 0.00 | 0.00 | 0.00 | 0.00 | 2.98 | 0.59 | 1.00 | 0.00 |
| His168Arg | 0.00 | 0.00 | 1.23 | 0.82 | 1.21 | 0.84 | 0.00 | 0.00 | 3.97 | 1.01 | 1.07 | 0.30 |
| Val143Ala | 0.00 | 0.00 | 0.00 | 0.00 | 0.00 | 0.00 | 0.00 | 0.00 | 3.92 | 0.31 | 1.16 | 0.40 |
| Ile195Thr | 0.00 | 0.00 | 0.00 | 0.00 | 0.00 | 0.00 | 0.00 | 0.00 | 2.00 | 0.00 | 2.07 | 0.89 |

The mutation Arg273His, which is the position 180 in the structural alignment, is a conservative mutation as both residues are polar positive according to the CINEMA color scheme. The Structure-based sequence alignment module shows that this column is highly conserved with Arg in approximately 89% of chains, His and Cys in approximately 5% each. The conservation on alignment position 180 is shown in Figure 2 from the *Additional file 1*. The accessibility, which is provided in Figure 6, is conserved but does not have very low values (ranges from 4 up to 39.7)(Table 3), as the column presents a light shade of blue. In regard to the topological properties (complex network metrics)(Table 3), shown in Figure 3 from the *Additional file 1*, the degree is conserved (3 up to 9); betweenness is not conserved as the column does not have a very similar shade; closeness is relatively conserved. Actually, in closeness, we see regions (a set) of conserved columns, which makes sense considering that if a vertex (residue) has a high closeness value, it is close to many vertices and it is likely that his neighbors present similar behavior. The same holds for vertices with low closeness values. Regarding the interactions established by column 180 (Table 4), the majority of residues in this position establish charged attractive, charged repulsive and hydrogen bonds, so these interactions are highly conserved. Hydrophobic interactions, provided in Figure 7, are not conserved, as there are only 8 chains (approximately 8%) that establish such interactions in this position. In Figure 4 from the *Additional file 1*, we show an example of how the domain specialist can inspect the specific interactions established by a residue at the atomic level. By clicking on any residue of the Interactions module, VERMONT shows the interactions established by a particular residue/atom in the context of protein 3D structure in a molecular viewer and in a 2D graph schematic representation. To sum up, we would not consider this mutation as damaging (which is in accordance with FoldX, which outlines this position with a gray rectangle) because the residue change is conservative, the accessibility is not low and there are few, non-conserved hydrophobic interactions in this position.

**Figure 6.**
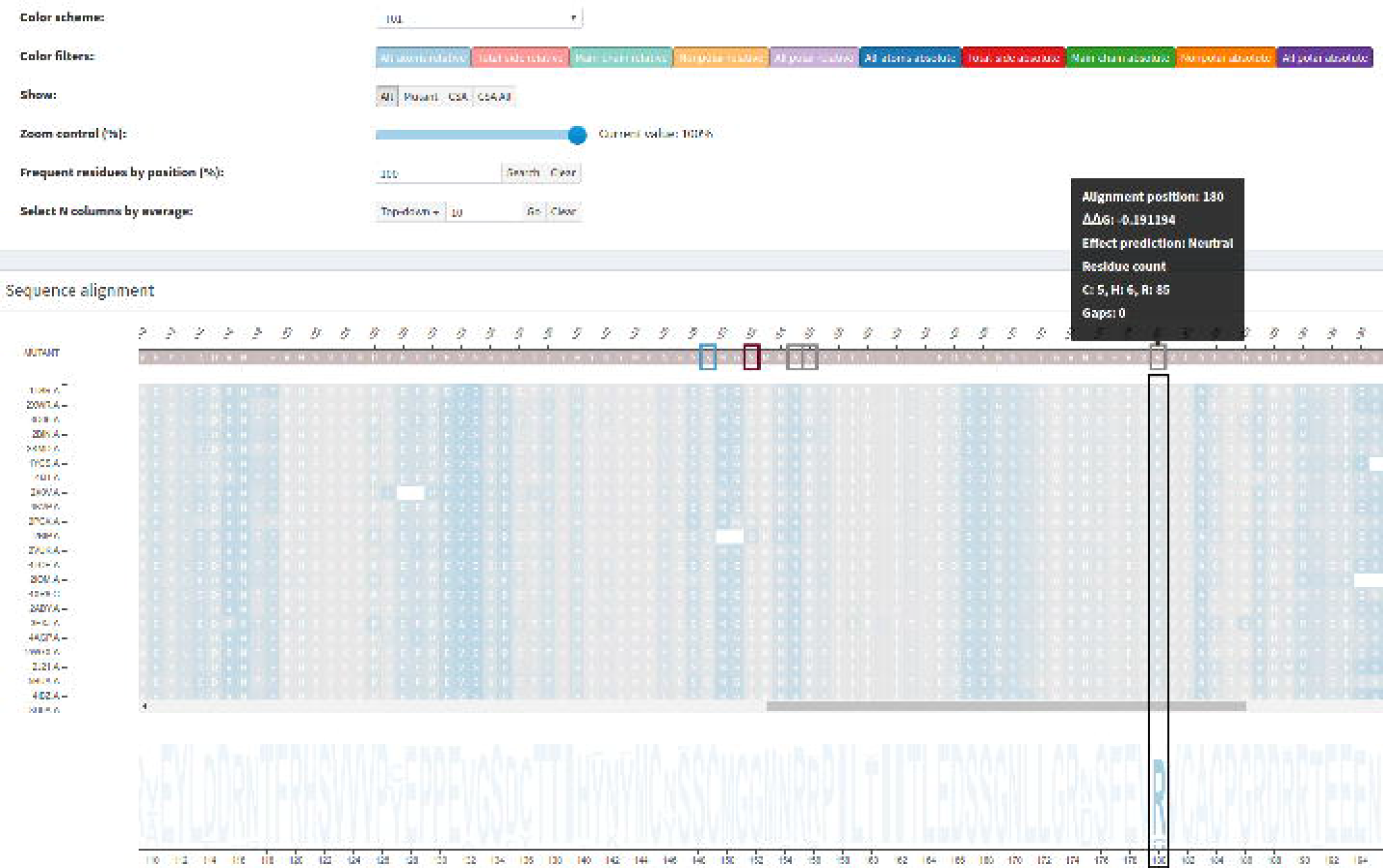
All-atoms relative accessibility. Low values and conservation highlighted for alignment position 180, which corresponds to mutation Arg273His in protein p53 (1TSR.A). The light shade of blue means that accessibility values are not very low.

**Figure 7.**
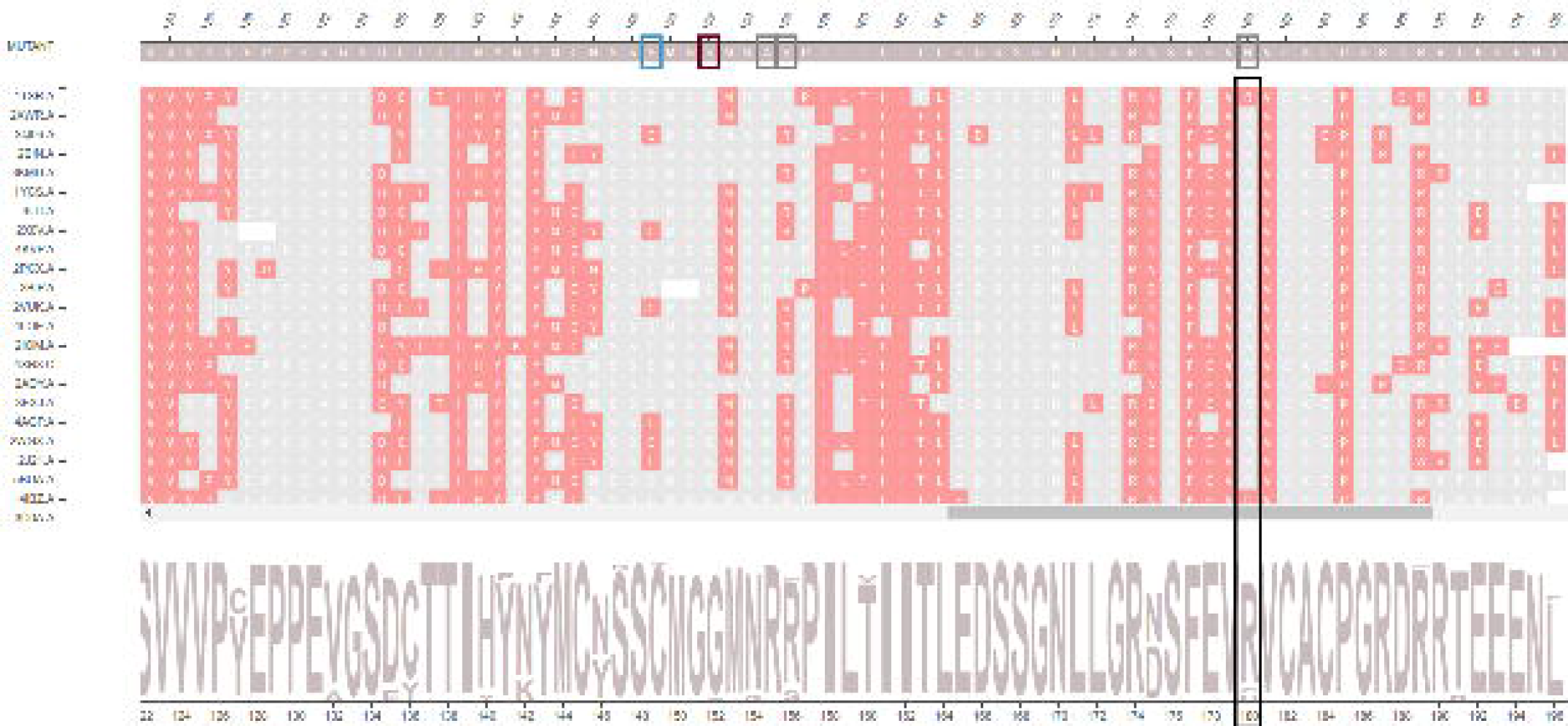
Hydrophobic interactions highlighted for alignment position 180. This position corresponds to mutation Arg273His in protein p53 (1TSR.A). Hydrophobic interactions are not conserved in this position as the majority of residues (approximately 92%) are not colored in rose.

Ile195Thr corresponds to position 102 in the structural alignment and is non-conservative, as Ile is non-polar aliphatic and Thr is polar neutral. Figure 8 shows column 102 is highly conserved, presenting only Ile residues. The accessibility in this column, provided in Figure 9 and Table 3, is low and conserved as the whole column presents a light shade of gray (0.3 up to 7.6). With regard to the topological properties (Table 3), shown in Figure 5 of the *Additional file 1*, the degree is relatively conserved (2 to 5); betweenness is not conserved and closeness is relatively conserved. In regard to the interactions in the alignment position 102, 100% of residues establish hydrogen bonds and 98% establish hydrophobic interactions (Table 4). Therefore, the hydrogen bonds and hydrophobic interactions are highly conserved. Figure 10 shows the conservation of hydrophobic interactions for alignment position 102. Figure 11 provides 3D and 2D views of interactions established by p53 wild type protein (1TSR.A, Ile195), showing that on the alignment position 102, the wild type protein presents only hydrogen bonds and hydrophobic interactions. Having these aspects in mind, we consider Ile195Thr as likely damaging, as it is non-conservative, with low and conserved accessibility and highly conserved hydrophobic interactions. This conclusion is in accordance with FoldeX, which outlines this position with a red rectangle.

**Figure 8.**
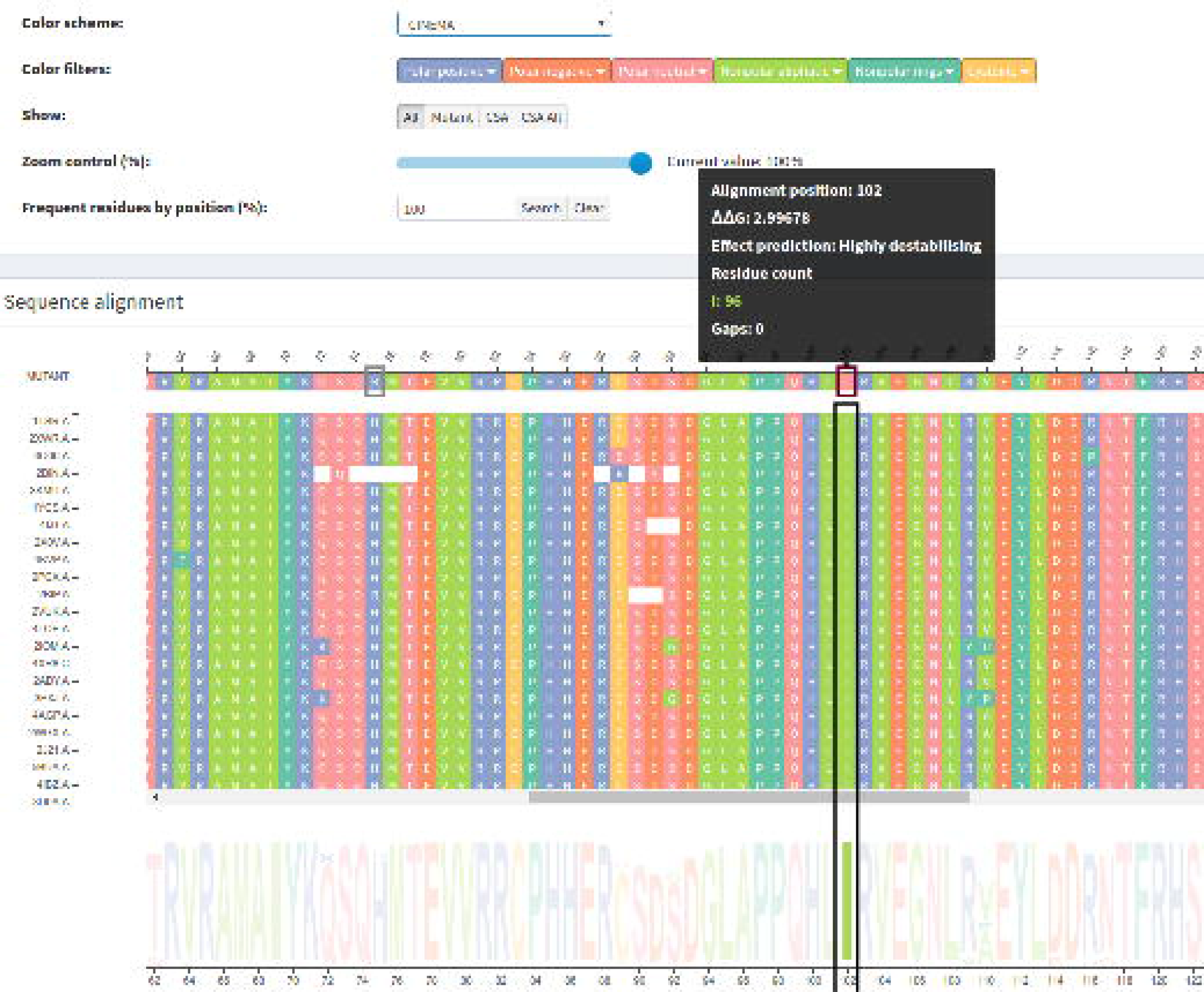
Residue conservation highlighted for alignment position 102. This position corresponds to the non-conservative mutation Ile195Thr in protein p53 (1TSR.A). This position is highly conserved, with only Ile residues, which can be seen in the sequence logo.

**Figure 9.**
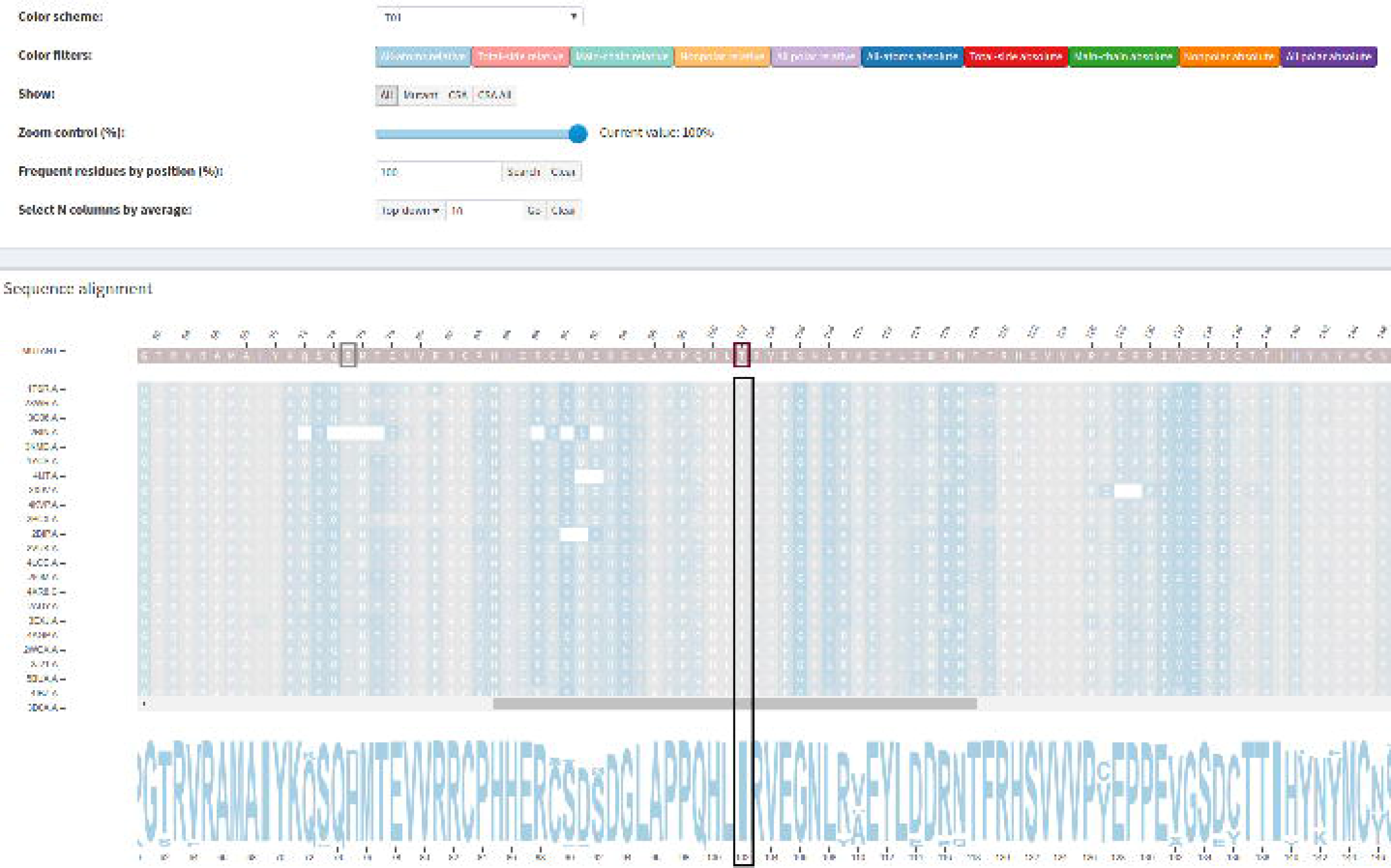
All-atoms relative accessibility highlighted for alignment position 102. This position corresponds to mutation Ile195Thr in protein p53 (1TSR.A). The light shade of gray in this whole column means that accessibility values are conserved and very low.

**Figure 10.**
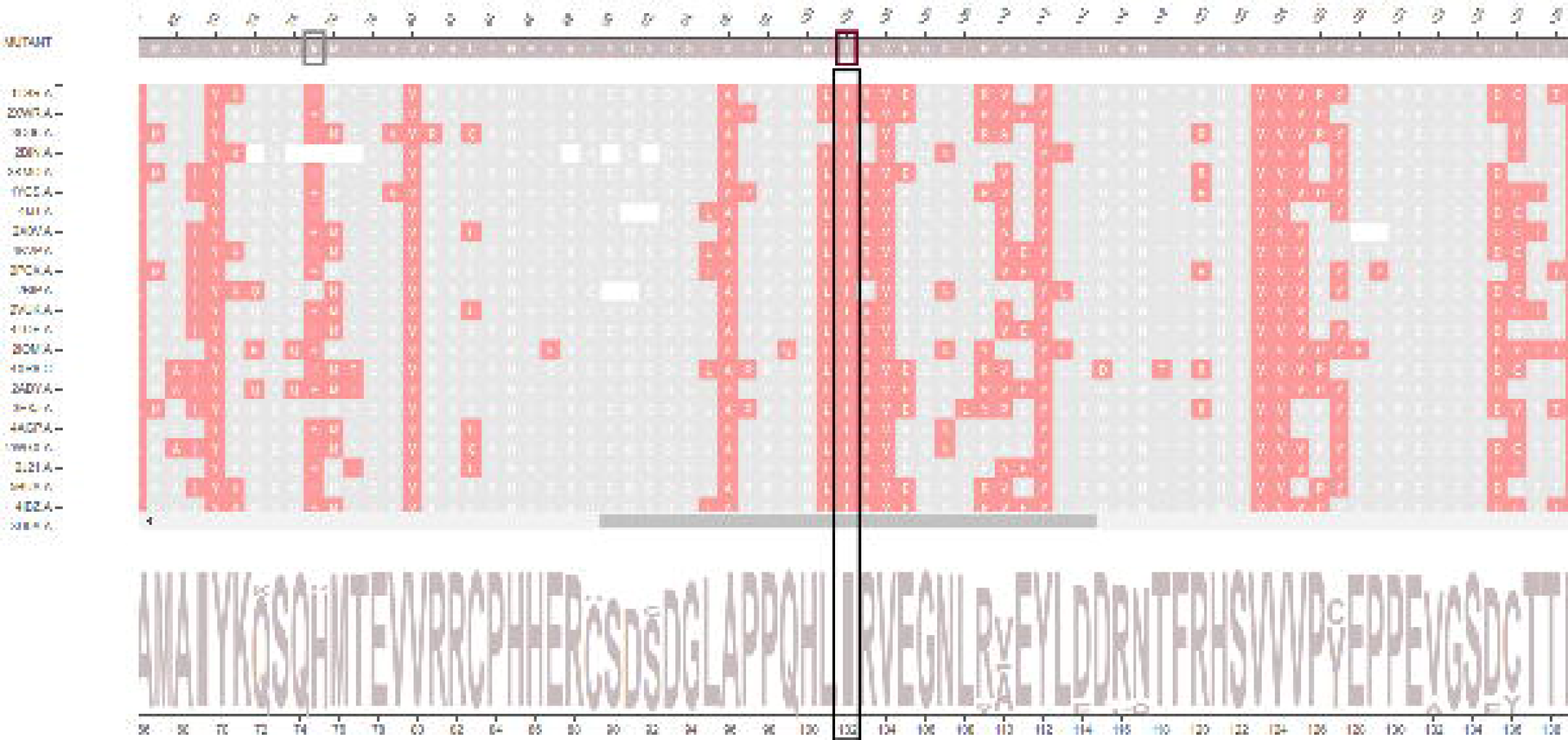
Hydrophobic interactions highlighted for alignment position 102. This position corresponds to mutation Ile195Thr in protein p53 (1TSR.A). Hydrophobic interactions are highly conserved in this position, as we see a column colored in rose.

**Figure 11.**
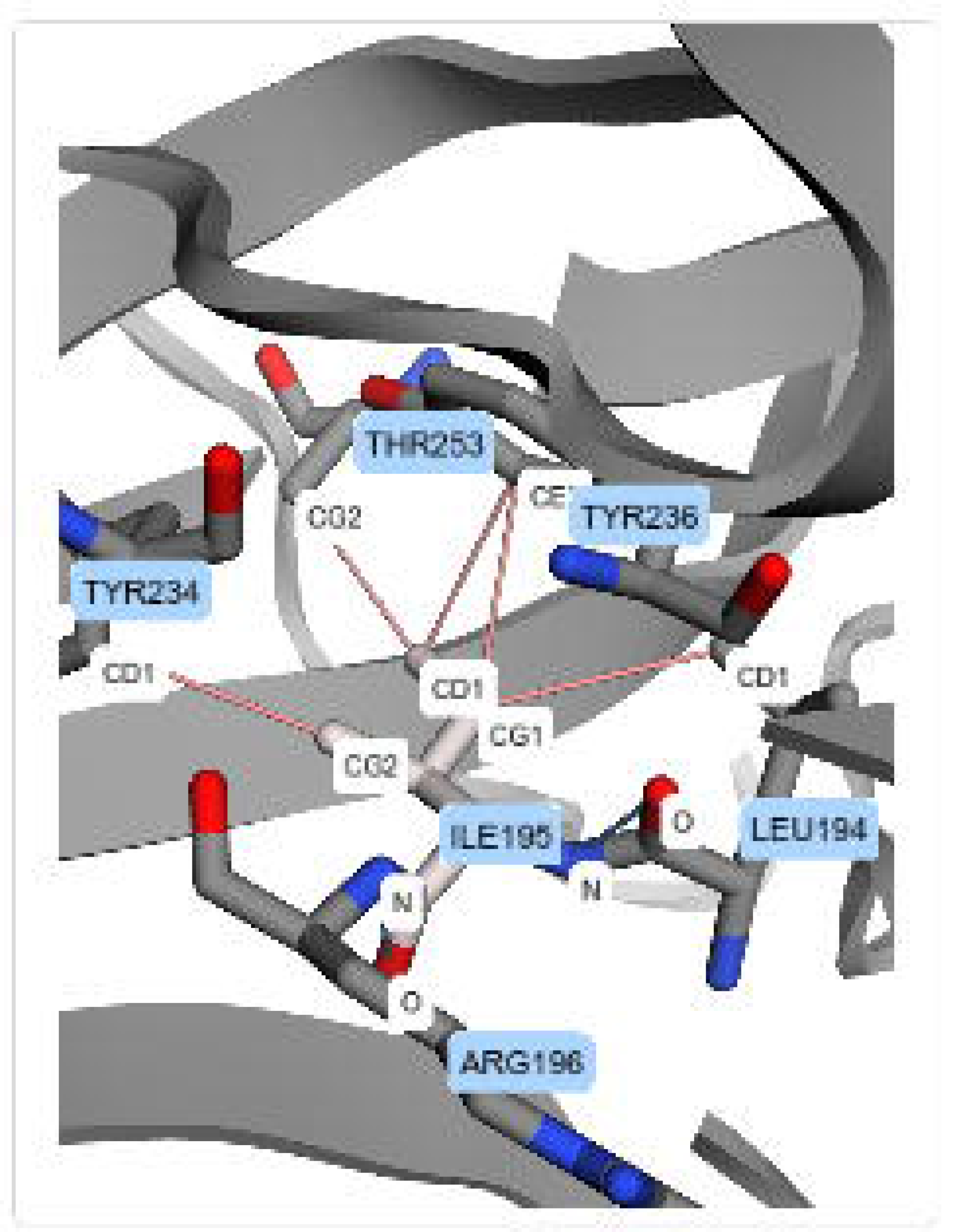
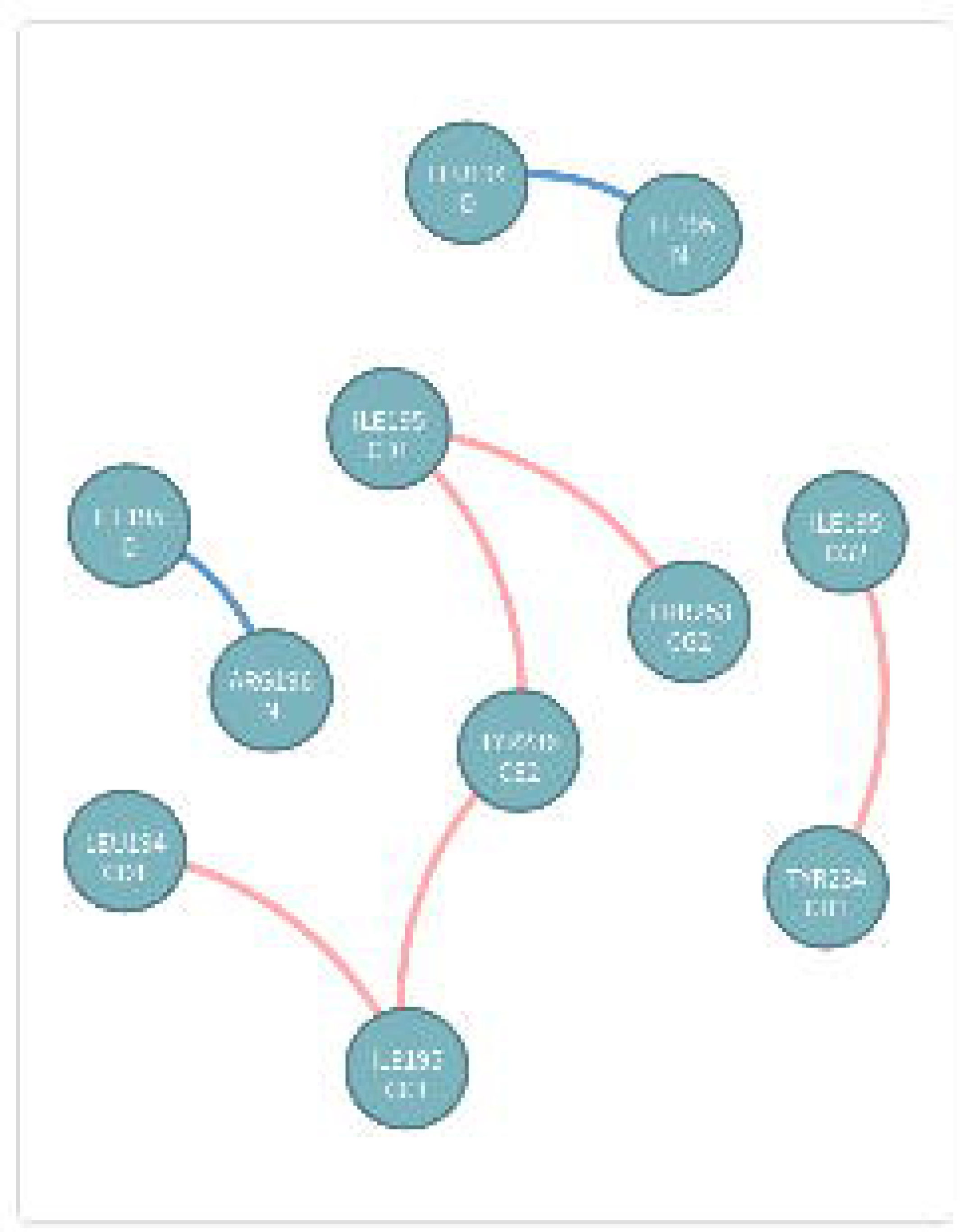
Interactions established by residue Ile195 of 1TSR.A (wild type protein). This residue is in the alignment position 102, which corresponds to mutation Ile195Thr in protein p53 (1TSR.A). Ile195 establishes hydrogen bonds and hydrophobic interactions. (A) 3D interactions for Ile195 of 1TSR.A in a molecule viewer. (B) 2D graph schematic representation for interactions from Ile195 of 1TSR.A

Analyzing these experimentally studied mutations through our visual platform and combining different structural and physicochemical data in a totally visual and interpretable manner, gives relevant clues that support the identification of damaging mutations. In the case of p53, it seems that mutations in positions with conserved and low accessibilities, coupled with conserved hydrophobic interactions, are likely to have an impact on protein structure/function. We could also analyze this dataset by looking for these specific characteristics to identify critical positions. For instance, the positions 145(Leu), 157 (Val) and 254 (Ile) in p53 (1TSR), which correspond to positions 52, 64 and 161 in the alignment, have very low and conserved accessibilities and have highly conserved hydrophobic interactions.

## Conclusions

This report presents VERMONT 2.0, a visual interactive platform that integrates sequence conservation, the intramolecular interaction network, solvent accessibility, physicochemical properties and complex network topological parameters, combining them with powerful interactive visualizations to make the impact of protein point mutations more understandable.

To assess the ability of VERMONT to gain insight into the impact of point mutations, we presented a use case in which we analyzed mutations that have been experimentally characterized. We show that VERMONT is able to identify these mutations in a completely visual manner, providing clues that help to identify those that potentially have an impact on structure/function. In this specific dataset, harmful mutations tend to present low and conserved values for accessibility, combined with conserved hydrophobic interactions.

As future work, we intend to design an automatic strategy to support users on the detection of harmful mutations, based on the structural and physicochemical properties computed by VERMONT. Additionally, we would like to investigate how VERMONT can be extended to address the multi-chain protein complex as a whole, as currently, we use individual chains. Last, we consider allowing domain specialists to use not only structures that are experimentally solved and available in PDB but also their own models.

## Abbreviations

ΔΔG: Gibbs free energy change, dTIM: defective triosephosphate isomerase, INPS: Impact of Non-synonymous variations on Protein Stability, HGMD: Human Gene Mutation Database, MAESTRO: multi-agent stability prediction upon point mutations, MD: Multi Descriptor, nsSNP: non-synonymous single-nucleotide polymorphisms, PDB: protein data bank, RINs: residue interaction networks, scTIM: saccharomyces cerevisiae triosephosphate isomerase, SDM: Site Directed Mutator, SNP: single-nucleotide polymorphisms, SVM: support vector machine, VERMONT: ViewER MutatiON Tool.

## Declarations

### Ethics approval and consent to participate

Not applicable.

### Consent for publication

Not applicable.

### Availability of data and material

Vermont interactive platform is available at:

http://bioinfo.dcc.ufmg.br/vermont/

## Authors’ contributions

SAS and RCM conceived the VERMONT platform. AVF and PMM designed and implemented the tool. SSG, SSA, and VSR implemented algorithms for property computation. AVF, SAS and RCM analyzed the results and wrote the manuscript. All authors read and approved the final manuscript.

## Acknowledgements

Not applicable.

## Additional Files

Additional file 1 — Supplementary material

Additional details and figures to support on the understanding and usage of VERMONT.

http://bioinfo.dcc.ufmg.br/vermont/download/supplementary-material.pdf

## References

1. International HapMap Consortium: The International HapMap Project. Nature 426(6968), 789–796 (2003). doi:10.1038/nature02168

2. Khafizov, K., et al.: Computational approaches to study the effects of small genomic variations. Journal of molecular modeling 21(10), 251 (2015)

3. Stenson, P.D., et al.: The human gene mutation database: towards a comprehensive repository of inherited mutation data for medical research, genetic diagnosis and next-generation sequencing studies. Human Genetics, 1–13 (2017)

4. Kulshreshtha, S., et al.: Computational approaches for predicting mutant protein stability. Journal of computer-aided molecular design 30(5), 401–412 (2016)

5. Worth, C.L., et al.: Sdm—a server for predicting effects of mutations on protein stability and malfunction. Nucleic acids research 39(suppl 2), 215–222 (2011)

6. Topham, C.M., et al.: Prediction of the stability of protein mutants based on structural environment-dependent amino acid substitution and propensity tables. Protein Engineering 10(1), 7–21 (1997)

7. Pires, D.E., et al.: mcsm: predicting the effects of mutations in proteins using graph-based signatures. Bioinformatics 30(3), 335–342 (2014)

8. Giollo, M., et al.: Neemo: a method using residue interaction networks to improve prediction of protein stability upon mutation. BMC genomics 15(4), 7 (2014)

9. Laimer, J., et al.: Maestro-multi agent stability prediction upon point mutations. BMC bioinformatics 16(1), 116 (2015)

10. Laimer, J., et al.: Maestroweb: a web server for structure-based protein stability prediction. Bioinformatics 32(9), 1414–1416 (2016)

11. Fariselli, P., et al.: Inps: predicting the impact of non-synonymous variations on protein stability from sequence. Bioinformatics 31(17), 2816–2821 (2015)

12. Savojardo, C., et al.: Inps-md: a web server to predict stability of protein variants from sequence and structure. Bioinformatics, 192 (2016)

13. Chen, C.-W., et al.: istable: off-the-shelf predictor integration for predicting protein stability changes. BMC bioinformatics 14(2), 5 (2013)

14. Capriotti, E., et al.: I-mutant2. 0: predicting stability changes upon mutation from the protein sequence or structure. Nucleic acids research 33(suppl 2), 306–310 (2005)

15. Cheng, J., Randall, A., Baldi, P.: Prediction of protein stability changes for single-site mutations using support vector machines. Proteins: Structure, Function, and Bioinformatics 62(4), 1125–1132 (2006)

16. Masso, M., Vaisman, I.I.: Accurate prediction of stability changes in protein mutants by combining machine learning with structure based computational mutagenesis. Bioinformatics 24(18), 2002–2009 (2008)

17. Dehouck, Y., et al.: Fast and accurate predictions of protein stability changes upon mutations using statistical potentials and neural networks: Popmusic-2.0. Bioinformatics 25(19), 2537–2543 (2009)

18. Parthiban, V., et al.: Cupsat: prediction of protein stability upon point mutations. Nucleic acids research 34(suppl 2), 239–242 (2006)

19. Pires, D.E., et al.: Duet: a server for predicting effects of mutations on protein stability using an integrated computational approach. Nucleic acids research, 411 (2014)

20. Shevade, S.K., et al.: Improvements to the smo algorithm for svm regression. IEEE transactions on neural networks 11(5), 1188–1193 (2000)

21. Silveira, S.A., et al.: Vermont: Visualizing mutations and their effects on protein physicochemical and topological property conservation. In: BMC Proceedings, vol. 8, p. 4 (2014). BioMed Central

22. Berman, H.M., et al.: The protein data bank. Nucleic acids research 28(1), 235–242 (2000)

23. Schymkowitz, J., et al.: The foldx web server: an online force field. Nucleic acids research 33(suppl 2), 382–388 (2005)

24. Okabe, A.: Spatial Tessellations: Concepts and Applications of Voronoi Diagrams, 2nd ed edn. Wiley series in probability and statistics. Wiley, Chichester; New York (2000)

25. Cgal, Computational Geometry Algorithms Library

26. Gonçalves-Almeida, V.M., et al.: Hydropace: understanding and predicting cross-inhibition in serine proteases through hydrophobic patch centroids. Bioinformatics 28(3), 342–349 (2012)

27. Santana, C.A., et al.: Gremlin: A graph mining strategy to infer protein-ligand interaction patterns. In: Bioinformatics and Bioengineering (BIBE), 2016 IEEE 16th International Conference On, pp. 28–35 (2016). IEEE

28. Sobolev, V., et al.: Automated analysis of interatomic contacts in proteins. Bioinformatics 15(4), 327–332 (1999)

29. Mancini, A.L., et al.: Sting contacts: a web-based application for identification and analysis of amino acid contacts within protein structure and across protein interfaces. Bioinformatics 20(13), 2145–2147 (2004)

30. Shatsky, M., et al.: A method for simultaneous alignment of multiple protein structures. Proteins: Structure, Function, and Bioinformatics 56(1), 143–156 (2004)

31. Kim, J., Wilhelm, T.: What is a complex graph? Physica A: Statistical Mechanics and its Applications 387(11), 2637–2652 (2008)

32. Csardi, G., Nepusz, T.: The igraph software package for complex network research. InterJournal Complex Systems, 1695 (2006)

33. Lee, B., Richards, F.M.: The interpretation of protein structures: estimation of static accessibility. Journal of molecular biology 55(3), 379–34004 (1971)

34. Ponnuswamy, P., et al.: Hydrophobic packing and spatial arrangement of amino acid residues in globular proteins. Biochimica et Biophysica Acta (BBA)-Protein Structure 623(2), 301–316 (1980)

35. Cheng, T.M., et al.: Prediction by graph theoretic measures of structural effects in proteins arising from non-synonymous single nucleotide polymorphisms. PLoS Comput Biol 4(7), 1000135 (2008)

36. Olivier, M., et al.: The iarc tp53 database: new online mutation analysis and recommendations to users. Human mutation 19(6), 607–614 (2002)

37. Friedler, A., et al.: Kinetic instability of p53 core domain mutants implications for rescue by small molecules. J. of Biological Chemistry 278(26), 24108–24112 (2003)

38. Joerger, A.C., et al.: Structures of p53 cancer mutants and mechanism of rescue by second-site suppressor mutations. Journal of Biological Chemistry 280(16), 16030–16037 (2005)

